# Temporal and regulatory dynamics of the inner ear transcriptome during development in mice

**DOI:** 10.1101/2022.10.17.512623

**Authors:** Rui Cao, Masaki Takechi, Xiuwan Wang, Toshiko Furutera, Taro Nojiri, Daisuke Koyabu, Jun Li

## Abstract

The inner ear controls hearing and balance, while the temporal molecular signatures and transcriptional regulatory dynamics underlying its development are still unclear. In this study, we investigated time-series transcriptome in the mouse inner ear from embryonic day 11.5 (E11.5) to postnatal day 7 (P7) using bulk RNA-Seq. A total of 10,822 differentially expressed genes were identified between pairwise stages. We identified nine significant temporal expression profiles using time-series expression analysis. The constantly down-regulated profiles throughout the development are related to DNA activity and neurosensory development, while the constantly up-regulated profiles are related to collagen and extracellular matrix. Further co-expression network analysis revealed that several hub genes, such as pnoc1, cd9, and krt27, are related to the neurosensory development, cell adhesion, and keratinization. We uncovered three important transcription regulatory paths during mice inner ear development. Transcription factors related to Hippo/TGFβ signaling induced decreased expressions of genes relate to the neurosensory and inner ear development, while a series of INF genes activated the expressions of genes in immunoregulation. In addition to deepening our understanding of the temporal and regulatory mechanisms of inner ear development, our transcriptomic data could fuel future multi-species comparative studies and elucidate the evolutionary trajectory of auditory development.

## Introduction

Hearing is the ability to receive and process sound through multiple organs. The inner ear plays a key role in obtaining hearing and balance (1, 2). Studying the temporal patterns in the inner ear transcriptome can unveil the dynamic genetic program underlying inner ear development. Given the unavailability of human samples due to ethical considerations, a mouse model is irreplaceable for studying expression and regulation of genes in the mammalian inner ear.

In mammals, the inner ear is derived from the otic placode, a thickened portion of the ectoderm, through a series of key stages (3). In mice, the otic placode invaginates through the otic cup and forms the otic vesicle at embryo development day 8 (E8) (4). These invaginate and form otic cups are eventually formed into otic vesicles/otocysts by embryo development day 9.5 (E9.5), the primordium of the future inner ear. At embryo development day 11.5 (E11.5), the otocyst elongates and forms cochlear pouch and vestibular pouch (5–7). At embryo development day 12.5 (E12.5), the utricle, saccule, and three semicircular canals of the vestibular organ are formed (8). The organ of Corti develops at the caudal wall of the cochlear duct at embryo development day 13 (E13), converting auditory stimuli into neural impulses after maturation (5, 6). Between embryo development day 16.5 (E16.5) and 18.5 (E18.5), hair cells differentiate and form a single row of inner hair cells and three rows of outer hair cells (6). Cellular patterns of hair cells complete and supporting cells present at postnatal day 0 (P0) (9). At postnatal day 7 (P7), the Corti and stria unscularis tunnels reach adult size; supporting cells form adult-like configurations (9).

Transcription factors (TFs) bind to specific DNA sequences to regulate transcriptional activities of the targeted genes (10), influencing otocyst induction, differentiation, patterning, morphogenesis and neurosensory development (11). Pathogenic mutations in several transcription factors regulating hearing-related genes are associated with deafness (12, 13), indicating key roles of the TFs in inner ear development. However, it is not fully understood how transcription factors and their regulatory gene network influence the development of inner ear over time.

A number of observational or interventional transcriptome studies have been conducted in the entire inner ear, specific tissues, or cell type at different stages of development (8, 14–17). To date, one comprehensive study conducted by Sajan et al. investigated the transcriptomic patterns of the overall inner ear and substructures (the presumptive cochlea, utricle, and saccule) from stage E9 to E15 using microarray (8). Some recent studies uncovered gene expression profiles in the mouse cochlea, vestibule (14, 15), hair cell (16) and spiral ganglion neurons (17) using RNA-Seq. The major limitations of these studies are 1) only two to five stages focused on a substructure of the inner ear have been included; and 2) differentially expressed genes or short time-series transcriptional patterns, instead of the regulatory gene network, were mostly studies.

To unveil the molecular basis and dynamic regulatory program during the inner ear development, we performed bulk RNA sequencing in the mouse inner ear from E11 to E18 with a one-day interval, as well in the postnatal stages, P0, P1, P3, and P7. We systematically investigated differentially expressed genes between stages, uncovered the temporal gene expression patterns, constructed co-expression networks, and inferred the temporal transcriptional regulatory network during inner ear development in mice.

## Result

### Differentially expressed genes and their temporal expression patterns

We first investigated the consistency/variability of the biological replicates based on the overall RNA expression profile. The result indicates that replicates from the same stage were overall clustered together according to Pearson’s correlation (Fig. 1b). To identify genes related to inner ear development, we performed differential expression analysis on 21,826 coding genes. We identified 10,822 differentially expressed genes (DEGs) (FDR < 0.001 and |log2 fold-change| > 1) between pairwise development stages (Table 1; Fig. 1c). The number of DEGs detected increases with the increased time interval between stages (Table 1). In addition, we identified four adjacent stage groups that presented relatively low differences in DEGs, including P0 and P1(40), E16.5 and E17.5 (117), E14.5 and E15.5 (171), and E12.5 and E13.5 (275), indicating relatively constant expression profiles across these periods.

**Table 1.** Pairwise different expression analysis

|  | B6E11 | B6E12 | B6E13 | B6E14 | B6E15 | B6E16 |
| --- | --- | --- | --- | --- | --- | --- |
| B6E11 | 0 | 865 | 1345 | 2339 | 3214 | 4168 |
| B6E12 | 865 | 0 | 275 | 1319 | 2065 | 3440 |
| B6E13 | 1345 | 275 | 0 | 855 | 1269 | 2947 |
| B6E14 | 2339 | 1319 | 855 | 0 | 171 | 1193 |
| B6E15 | 3214 | 2065 | 1269 | 171 | 0 | 941 |
| B6E16 | 4168 | 3440 | 2947 | 1193 | 941 | 0 |
| B6E17 | 4754 | 3938 | 3075 | 1543 | 1011 | 117 |
| B6E18 | 4324 | 4133 | 3521 | 1944 | 2045 | 274 |
| B6P0 | 4520 | 4461 | 3708 | 2169 | 1912 | 669 |
| B6P1 | 4836 | 4843 | 4116 | 2608 | 2373 | 939 |
| B6P3 | 5013 | 4743 | 4243 | 2625 | 2484 | 660 |
| B6P7 | 5760 | 5516 | 5032 | 3809 | 3540 | 1904 |

| B6E17 | B6E18 | B6P0 | B6P1 | B6P3 | B6P7 |
| --- | --- | --- | --- | --- | --- |
| 4754 | 4324 | 4520 | 4836 | 5013 | 5760 |
| 3938 | 4133 | 4461 | 4843 | 4743 | 5516 |
| 3075 | 3521 | 3708 | 4116 | 4243 | 5032 |
| 1543 | 1944 | 2169 | 2608 | 2625 | 3809 |
| 1011 | 2045 | 1912 | 2373 | 2484 | 3540 |
| 117 | 274 | 669 | 939 | 660 | 1904 |
| 0 | 339 | 304 | 679 | 707 | 1944 |
| 339 | 0 | 441 | 528 | 261 | 1430 |
| 304 | 441 | 0 | 40 | 473 | 1382 |
| 679 | 528 | 40 | 0 | 395 | 1450 |
| 707 | 261 | 473 | 395 | 0 | 657 |
| 1944 | 1430 | 1382 | 1450 | 657 | 0 |

**Figure 1.**
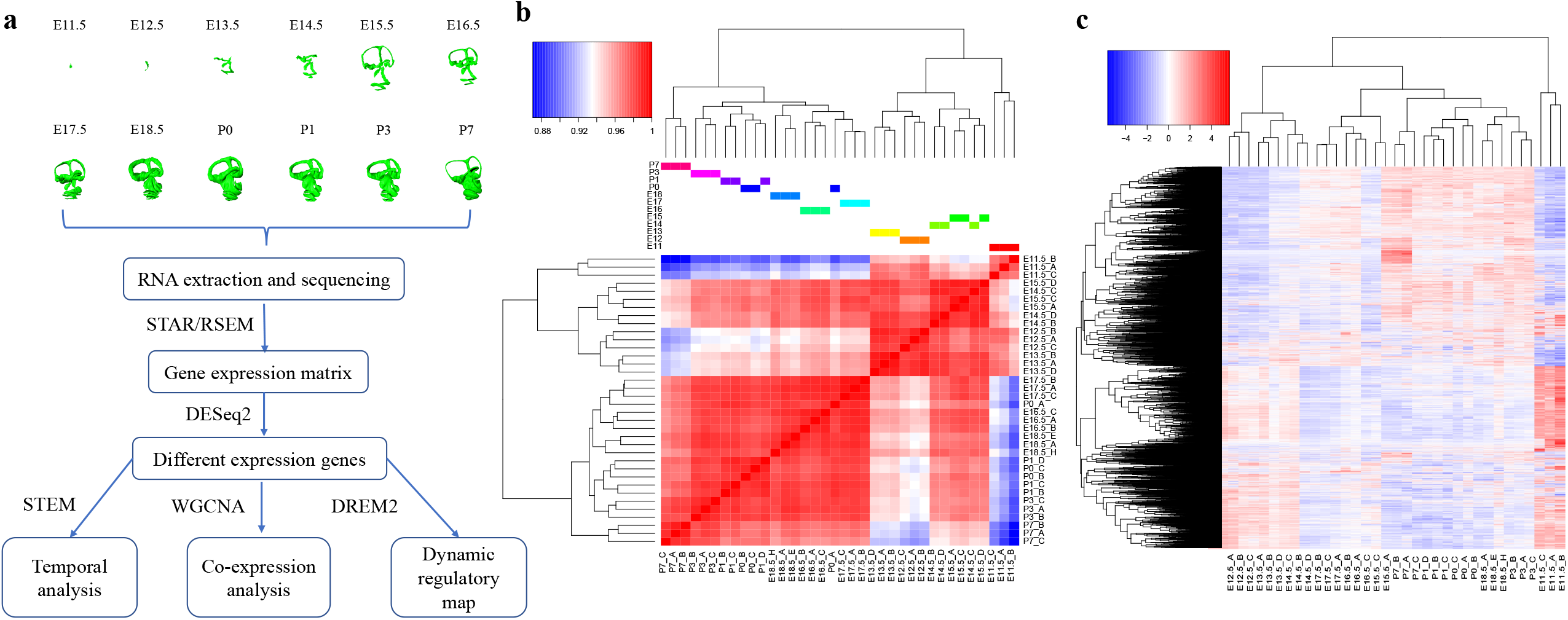
Study design and Global gene expression profile. (a) The study design and analytical pipeline; (b) Pearson correlation between samples based on the overall expression profile; (c) Hierarchical clustering of DEGs and samples (FDR < 0.001 and |log2 fold-change| > 1).

To disclose the dynamic expression patterns of DEGs at different developmental stages, we grouped 10,822 DEGs into 50 different temporal expression profiles using short time-series expression miner (STEM). A total of 7,207 DEGs were clustered into 9 profiles (p-values ≤0.05) (Fig 2). Temporal expression profiles 39, 25, and 19 showed overall decreased expression from E11.5 to P7. Genes within these three profiles were enriched in the GO annotations related to the neuron system, inner ear development, and DNA replication, such as “DNA repair” (GO:0006281), “axonogenesis” (GO:0007409), and “inner ear development” (GO:0048839) (Supplementary Data S1). In contrast, profiles 43, 44, 26, and 36 showed an overall increased expression from E11.5 to P7, mostly enriched in extracellular matrix organization and collagen matrix, such as “extracellular matrix organization” (GO:0030198) and “collagen metabolic process” (GO:0032963), and in immunological terms, such as “leukocyte mediated immunity” (GO:0002704) and “regulation of immune effector process” (GO:0002698) (Supplementary Data S2). Profiles 4 and 49 exhibited a biphasic expression pattern with decreased expressions at early stage while increased the expression afterwards, and them upregulate, uniquely enriched in the cell activity, such as “cell-substrate adhesion” (GO:0031589) (Supplementary Data S3). Overall, the temporal expression analysis revealed the distinct and dynamic expression patterns of DEGs over the course of development. Genes related to collagen-containing extracellular matrix and immunoregulation presented increased expression during development, while genes related to inner ear development and DNA replication revealed decreased expression during the development.

**Figure 2.**
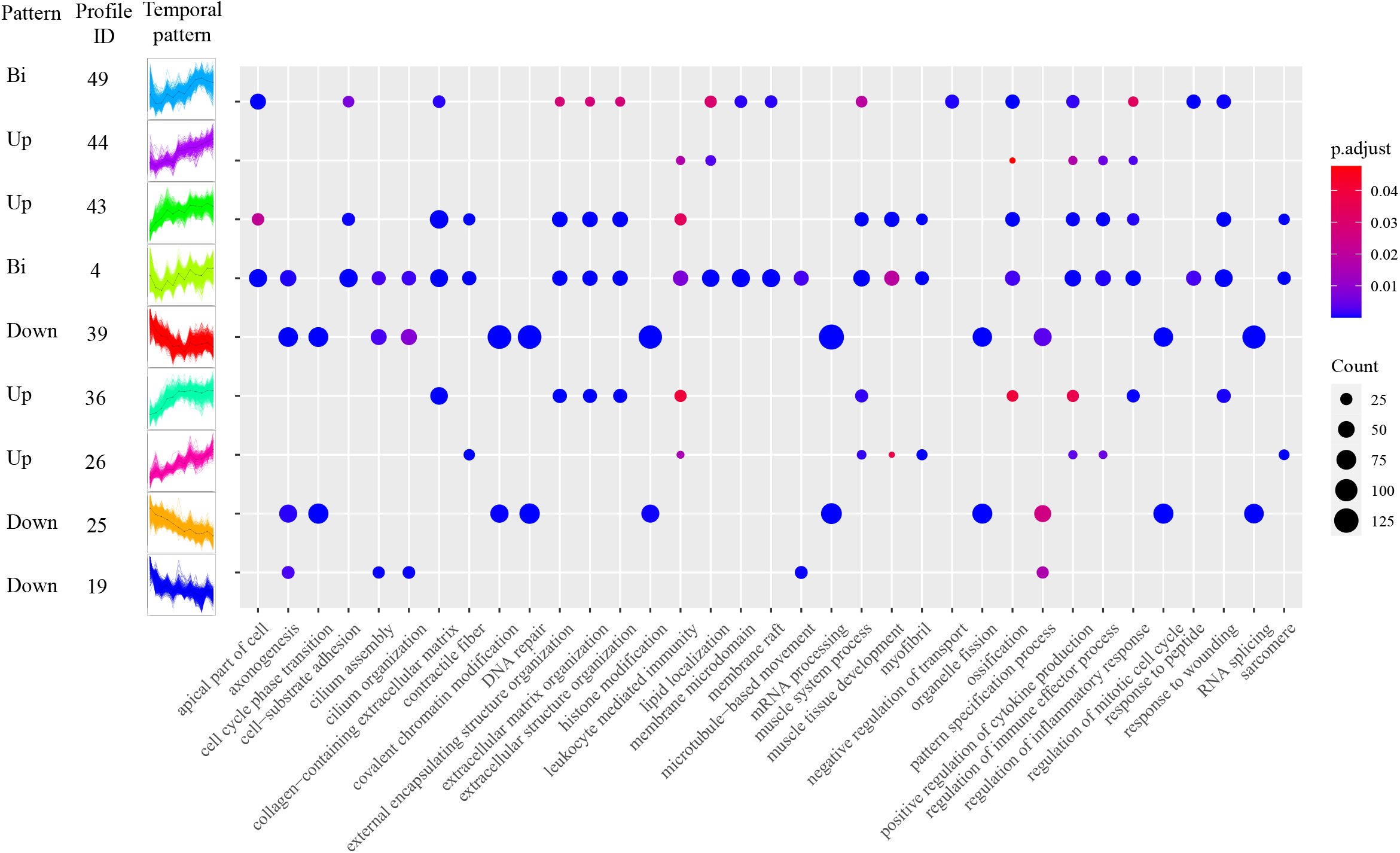
Temporal expression patterns of all the DEGs and their GO enrichment result. The left panel showed nine statistically significant Profiles (P<0.05) temporal expression pattern, assigned into three expression pattern, namely Bi, Up, Down. The right panel showed each profile GO enrichment result. Full enrichment result can be found in Supplementary Data S1-S3.

### Temporal expression patterns of genes related to hearing

We investigated the temporal patterns of 407 DEGs associated with hearing. A cluster of 198 genes exhibited relatively high expression in E11.5, then decreased and maintained at a relatively low expression level after E14.5. Another cluster of 207 genes showed increased expression over the development stages (Fig 3). Unsurprisingly, “inner ear development” is the common enriched GO term between the two clusters. However, the two clusters present unique enriched GO terms in molecular function and cellular component. For example, the down-regulated cluster is related to the Wnt pathway, such as “Wnt signaling pathway” (GO:0016055) and “Wnt-protein binding” (GO:0017147), which relate to early inner ear development, such as otic specification (Fig 3c) (18). In contrast, the up-regulated cluster is mainly enriched in “collagen-containing extracellular matrix” and “ion channel activity,” which are vitally important to transduce sounds into electrical signals in cochlea (Fig 3e) (19).

**Figure 3.**
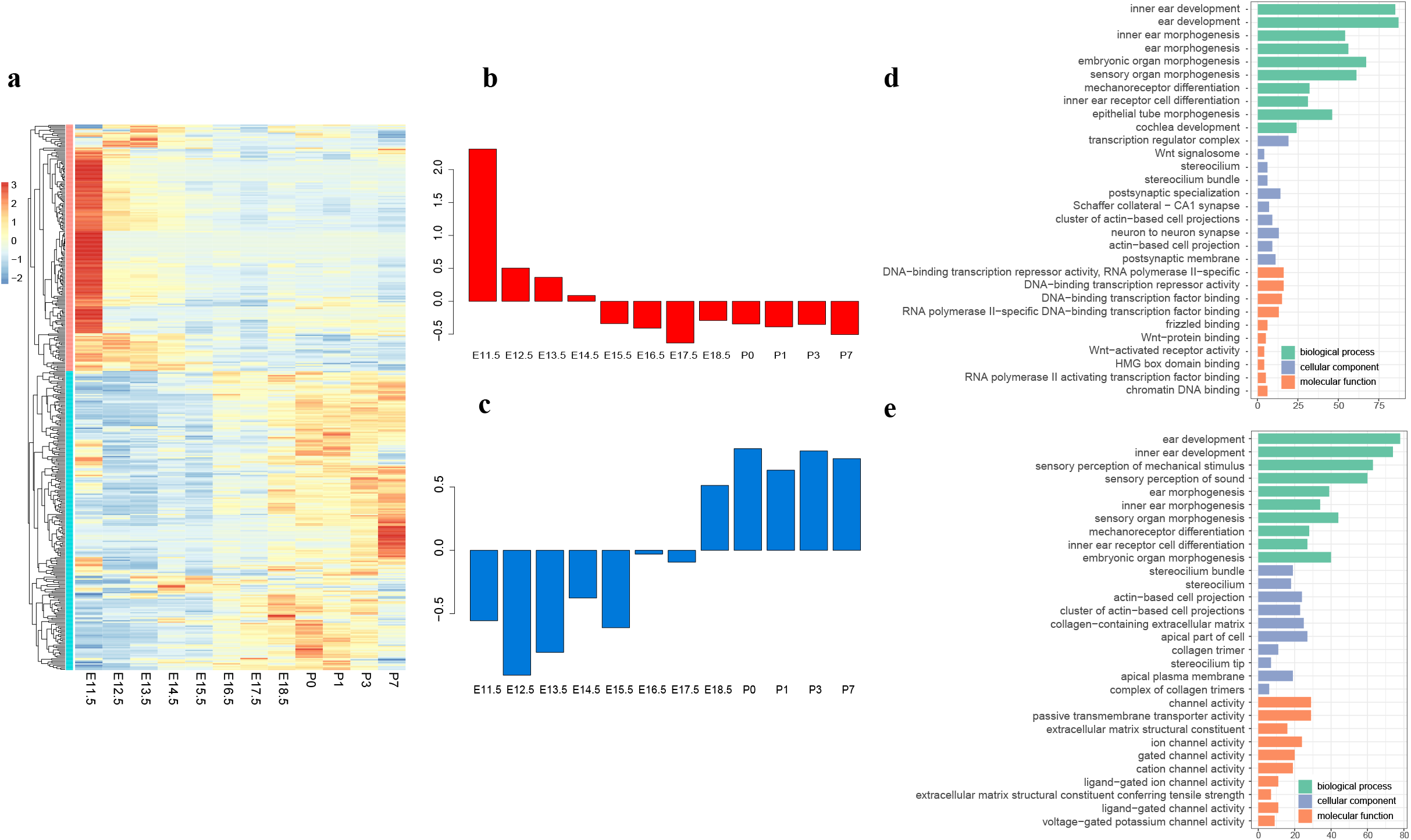
Expression of hearing related genes and the pathways or biological processed they are involved. (a) Expression of hearing related genes; (b) Downregulated cluster of genes and their average temporal expression levels; (c) Upregulated cluster of genes and their relative expression value; (d) Top 10 enrichment GO terms for downregulated cluster; (e) Top 10 enrichment GO terms for upregulated genes. Details of the enrichment result for hearing related genes can be found in Supplementary Data S4.

### Co-expression network analysis

We adopted weighted gene co-expression network analysis (WGCNA) to disclose the co-expression patterns among 10,882 DEGs. As a result, a gene co-expression network with 19 modules was constructed. The largest module, Turquoise, contains 5,112 genes. Another five largest modules included the Blue (1,837 genes), Brown (1,804 genes), Yellow (477 genes), Green (267 genes), and Red (266 genes) modules (Supplementary Fig S1).

To obtain functional explanations for each module, we performed enrichment analyses in GO terms and KEGG pathways. Genes in the Brown and Blue modules are significantly enriched in GO terms related to collagen and extracellular matrix, such as “collagen metabolic process” (GO:0032963), “collagen-containing extracellular matrix” (GO:0062023), and “extracellular matrix binding” (GO:0050840) (p-values <0.001, Fisher’s exact test). The KEGG pathways in the Brown and Blue modules were enriched in “ECM-receptor interaction” (mmu04512), “cytokine-cytokine receptor interaction” (mmu04060), and “cell adhesion molecule” (mmu04514) (p-values <0.001, Fisher’s exact test; Supplementary Data S6-S7). Genes in the Turquoise module were enriched in DNA activity and neuron system development, such as “DNA replication” (GO:0006260), “DNA packaging” (GO:0006323), “DNA recombination” (GO:0006310), “axonogenesis” (GO:0007409), “inner ear development” (GO:0048839), and “sensory organ morphogenesis” (GO:0090596) (p-values <0.001, Fisher’s exact test). Several important KEGG pathways related to inner ear development were enriched in the Turquoise module, such as “Wnt signaling pathway” (mmu04310) and “Notch signaling pathway” (mmu04330) (p-values <0.001, Fisher’s exact test) (Supplementary Data S5). Overall, the WGCNA analysis revealed three functionally important co-expression modules. The functionality these three modules revealed here is overall consistent with temporal expression patterns revealed by STEM analysis–increased expression of genes related to extracellular matrix organization and decreased expression of genes related to inner ear development and DNA replication.

We further identified hub genes with the highest degree centrality from the three largest co-expression modules (Turquoise, Blue and Brown) (Fig 4), which are biologically relevant to the mouse inner ear development. In the Turquoise module, the most connected gene, Krt27, encodes type I keratin. The other genes with highest connectivity are also mostly keratin-coding genes, such as Krt26, Krt33b, Krt71, Krt31, and Krt32 (Supplementary Table S2). In the Blue module, the most connected gene is Cd9 (Supplementary Table S2) that encodes a member transmembrane 4 superfamily and is involved in multiple biological processes, such as adhesion, motility, membrane fusion, signaling and protein trafficking (20). In the Brown module, the gene with the highest connectivity is Pnoc (Prepronociceptin), which is involved in chemical synaptic transmission. Among top ten genes with the highest connectivity in the module Turquoise, eight genes, including Pnoc, Nckap5, Zswim5, Dpysl5, Dner, Rnf165, Slc1a2, and Crmp1, are related to the neuron system development and regulation (Supplementary Table S2). In summary, the functionality of the hub genes from three largest co-expression modules indicated their key roles in the neurosensory and development of the inner ear.

**Figure 4.**
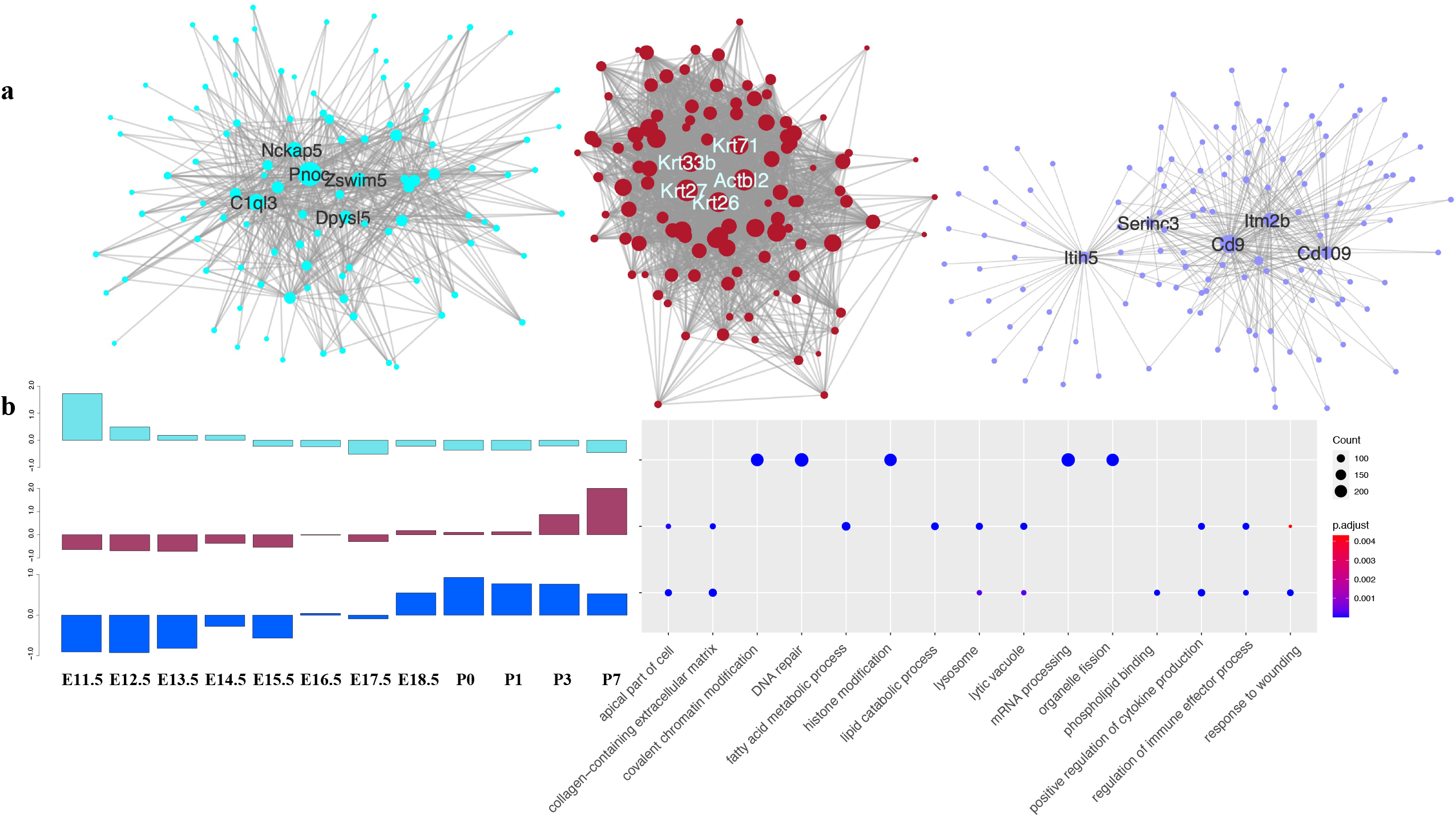
The network structure, temporal expression patterns, and enriched GO terms of top co-expression modules. (a) Three networks from Module Turquoise, brown and Blue visualized by Cytoscape. The circle size was mapped based on edge numbers. Top five genes with highest connectivity were labelled. (b) Relative gene expression and their GO enrichment result. The left panel is the relative gene expression of each module. The TPM value were Z-score normalization and calculate average expression per stage; the right figure is the enrichment GO terms for each module.

### Dynamic regulatory networks

To reveal the transcriptional regulation events related to inner ear development in mice, we reconstructed dynamic regulatory networks using DREM2 (21) based on our time-series RNA-Seq data. A total of 7,727 genes with predicted TF-gene binding relationships were assigned to 39 regulatory paths containing a series of bifurcation events, which occur when a group of genes simultaneously level up or repress the expression level (21) (Fig 5a). We identified three major paths regulated by six transcription factor sets (Fig 5b-d and Supplementary Data S9). Path A was regulated in E11.5 and E12.5 by TFs from the Sox and E2F gene family, such as Sox3, Sox2, and E2F2 (Fig 5b and Supplementary Data S9). The regulatory events in Path B occurred in E12.5 and E15.5, activated by several TFs, such as E2F, GATA, STAT and SP (Fig 5c and Data S9). The mostly enriched GO terms of the regulated genes in Path A and B are both “axonogenesis” (GO:0007409) (Fig 5b, c and f). In addition, GO terms “inner ear development” (GO:0048839) and “inner ear morphogenesis” (GO:0042472) are both enriched for the regulated genes in path A and B, while the related TFs are enriched in pathways related to inner ear development, such as TGF-beta signaling pathway and Hippo signaling pathway (Fig 5e). These results indicated that multiple and parallel regulatory processes have been involved in the inner ear development. We further discovered that the up-regulate path C was upregulated in E12.5 and E15.5 by five interferon regulatory factor (IRF) IRF3, IRF4, IRF5, IRF8, IRF9 (Fig 5d and Supplementary Data S9), and the transcription factor were enriched in several immune response pathway, such as human T-cell leukemia virus 1 infection, Toll-like receptor signaling pathway (Fig 5e, Supplementary Data S8). As expected, the enriched GO terms of activated genes in path C are mainly related to immunoregulation, such as “regulation of immune effector process” (GO:0002697) and “regulation of inflammatory response” (GO:0050727) (Fig 5d and f).

**Figure 5.**
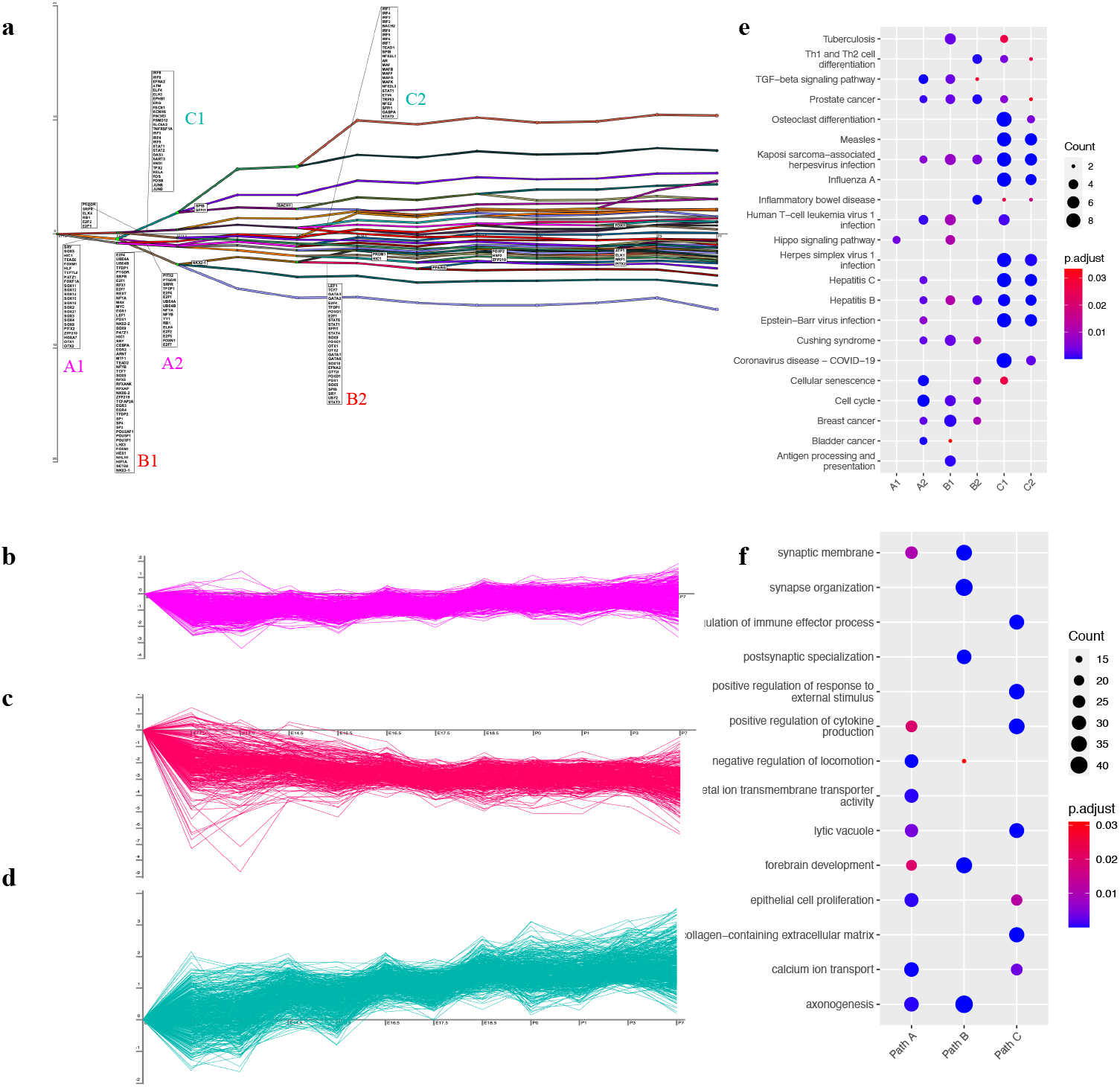
Temporal transcriptional regulatory map using DREM2. (a) Major dynamic regulatory paths during inner ear development. The x-axis denotes time-points, the y-axis denotes the gene expression level that relative to E11.5. Six key TF groups labelled as A1, A2, B1, B2, C1, C2; (b) Temporal expression pattern of TFs targeted genes of path A. (c) Temporal expression pattern of TFs targeted genes of path B. (d) Temporal expression pattern of TFs targeted genes of path C. (e) Enriched KEGG pathways for TFs. Full enrichment result can be found in Supplementary Data S10. (f) Enriched GO terms for gene in path A, B, C. Full enrichment result can be found in Supplementary Data S9

## Discussion

The inner ear is an essential component for sensing sound. Understanding the molecular mechanism of normal inner ear development in mammals is crucial for future therapy of hearing disorders. We analyzed the temporal expression of 21,826 coding genes in the mouse inner ear during prenatal and postnatal periods using bulk RNA-Seq. We identified distinct temporal gene expression clusters, revealed several co-expression modules, and discovered key transcriptional regulatory processes related to the hearing development. To the best of our knowledge, this is the first temporal transcriptomic study of inner ear development in mice with dense sampling covering both prenatal and postnatal periods to reveal the gene regulatory processes underpinning inner ear development. In addition, our high-throughput transcriptomic NGS data could benefit future comparative transcriptomic studies of inner ear development across mammals, potentiating studies on the evolutionary dynamics of auditory development at the molecular level.

Our co-expression network and time-series expression analyses revealed that genes highly expressed in the early embryo stage and repressed expression at later stages are related to the neuron system development (Fig. S1; Supplementary Data S1). Eight out of ten hub genes in the co-expression module Turquoise are involved in neurosensory development. The Pnoc gene, mostly connected in the co-expression network, is a synaptic adaptor protein involved in excitatory synaptic transmission. The Pnoc gene binds to the nociceptin receptor to increase pain sensitivity (22). The expression of the Pnoc gene is mainly detected in the central nervous system (23) and mutations in this gene could lead to abnormal mechanical nociception (24). In our study, the high expression of the Pnoc gene may contribute to the early development of neurosensory.

Our results revealed that several enriched GO terms of upregulated DEGs are related to collagen and extracellular matrix, such as “collagen-containing extracellular matrix”, during postnatal periods. The extracellular matrix plays an essential role when sounds are received and transformed into electrical signals (25). Collagens are major constituents of the extracellular matrix. Specific mutations in several collagen genes have been reported to be responsible for hearing disorders, such as Stickler syndrome (26) and Alport syndrome (27). Our results indicated that the collagen genes were upregulated in the postnatal stages, especially at P3 and P7 (Fig 2). Many collagen genes, such as Col1a1, Col2a1, and Col3a1, show a pattern of increased expression over time and peak at P3 and P7. Consequently, high expression levels of the collagen genes might result in collagen formation, maintaining normal function to perceive sound in the cochlea.

One interesting finding in our study is that several keratin hub genes (such as Krt27 and Krt26) in co-expression module Brown present high expression levels in the stage P7, indicating their critical influence on the inner ear development at a later stage. Keratin is one of the structural fibrous proteins, the key structural material making up hair, nails, feathers, horns, claws, hooves, and the outer layer of skin in vertebrates (28). Keratin was reported to be localized in the cochlear duct in the inner ear of mice (29). The expression of multiple genes from keratin family might synergistically contribute to the development of cochlear structure, but functional contribution of specific keratins to cochlear development is still unclear.

We employed DREM2 to infer transcriptional regulatory network during the inner ear development. Most regulatory events occurred between stage E11.5 and E13.5, indicating this period is pivotal for the transcriptional regulation during the inner ear development. We identified two downregulated paths A and B that are enriched in neurosensory and inner ear development. The enriched KEGG pathways of TFs in these two paths are related to Hippo signaling and TGFβ signaling (Fig 5e), which play important role in mouse inner ear development (30–36). In detail, the Hippo signaling pathway can maintain organ size and homeostasis by regulating cell proliferation, differentiation, and apoptosis (31). *In vitro* experiments showed that the upregulated Hippo signaling pathway reduced hair cell loss, while downregulated Hippo signaling pathway increased hair cell vulnerability in the inner ear (33). The TGFβ signal transduction pathway can mediate the expression of retinoic acid (RA), which regulates cell differentiation and normal patterning (35). Either an excess or deficiency of RA occurred at a critical stage can result in a malformation of the inner ear morphology (35). One recent study demonstrated that TGFβ signaling and insulin-like growth factor type 1 (IGF-1) jointly coordinated the cellular dynamics required for morphogenesis of the mouse inner ear, supporting the critical role of TGFβ signaling in inner ear cell senescence (34). Neuronsensory downregulated by Hippo signaling and TGFβ signaling may relate to prosensory identity. Previous studies has identified the progressively loss of prosensory domain in cochlea, which is initial in the whole cochlea and final degenerate to certain prosensory domain and the lateral region of Kölliker’s organ during development (5).Transcript factor Sox2, which identified in our regulate map, play important role during this process (37, 38). Therefore, it is unsurprised that overall gene expression related to neurosystem development will decrease. Taken together, our study supports the key regulatory roles of Hippo signaling and TGFβ signaling pathways in the cell differentiation and neurosensory development of the mice inner ear.

Gene regulatory path C in our study was activated by a series of IRFs in E12.5 and E15.5 (Fig 5d and e), enriched with GO terms related to immune responses. Previous studies demonstrated that the presence of certain immune cells present in the mature inner ear is linked to the hair cell damage in both the cochlea and vestibular organs of mammals (39–41). The up-regulation of certain immunoregulatory pathways starting from E12.5 may be related to the immune cell proliferation or apoptosis in the developmental process, which has been documented beginning from the otic cup stage to birth (42–44). Since immune-mediated inner ear disease and sensorineural hearing loss have been widely recognized (45, 46), further studies of TFs related to immunoregulation may shed light on the future immunotherapy for the inner ear diseases.

A handful of similar studies have been conducted to investigate the gene expression profile of the mouse inner ear (8, 14–17). Among those studies, Sajan et al. focused on the developmental transcriptome of the mouse inner ear in the period of E9-E15 (8), which overlapped several stages of our study. Sajan’s study revealed 28 different expression patterns between time points or tissues and identified 50 pathway cascades. The marked differences between our study and Sajan’s study reside in 1) developmental stages covered; and 2) sequencing and data mining techniques used; and 3) whether gene regulatory processes were discovered. RNA-Seq-based studies provide more accurate and higher sensitivity results than the microarray technique (47), which was deployed in the study of Sajan et.al. In addition, we quantified the expression of 21,826 coding genes, which is much greater than the number (14,065) in Sajan’s study. We also incorporated TF-gene interaction information and identified three regulatory paths that could influence mouse inner ear development, and this key analysis was omitted in Sajan’s study. The expanded candidate gene set, co-expression network analysis, temporal expression analysis and gene regulatory network reconstruction allow more comprehensive expression profiling and pattern recognition to unveil biologically relevant signatures for inner ear development across different stages.

The main limitation of our study is that no specific tissues, such as the vestibular organ or cochlea, were included. Therefore, although we identified several transcriptional signatures related to cochlear development, no transcriptional signature of neuron differentiation or hair cell development has been discovered in the late embryo or postnatal stages. Future studies may consider incorporating high-density postnatal stages to discover subtle transcriptional regulatory signatures in the postnatal stages of inner ear development.

## Conclusion

We provided a comprehensive temporal transcriptomic analysis to unveil the molecular basis of inner ear development in mice. We quantified the gene expression profile during the key developmental stages of inner ear development, which spans both the prenatal and postnatal periods. DEGs related to neurosensory development and collagen synthesis have been identified. The temporal and co-expression analysis revealed two important temporal expression patterns: the downregulated gene cluster is enriched in the development of neurosensory and inner ear, while the upregulated gene cluster is enriched in cochlea structural formation. The reconstructed gene regulatory network showed that the downregulated TFs in Hippo signaling and TGFβ signaling transcriptionally regulated genes related to the neurosensory and inner ear development between stages E11.5 and E15.5. Our study unveiled the temporal and regulatory transcriptomic dynamics during the inner ear development, paving the way for future comparative transcriptomic studies to elucidate the evolutionary trajectory of mammal hearing.

## Methods

### Animals

Ethics permits to conduct animal experiments have been assessed and approved at Tokyo Medical and Dental University, Japan (A2019-060C3 and A2021-198A). Experiments were performed in accordance with the Japanese law on animal welfare and carried out in compliance with the ARRIVE guidelines. We collected 36 embryos and newborn C57/BL6 mice bred at Tokyo Medical and Dental University. Mice were sacrificed by cervical dislocation. Considering that primordia of the inner ear start to form around or after E10, we studied prenatal stages from E11 to E18 with one-day intervals. We also included four time points from P0 to P7 when the tunnel of Corti reached adult size (9). In total, twelve developmental stages were identified and labeled: E11.5, E12.5, E13.5, E14.5, E15.5, E16.5, E17.5, E18.5, P0, P1, P3, and P7 (Fig 1a). Each stage had three replicates.

### RNA extraction and sequencing

The inner ear regions of the mouse embryo were carefully clipped with a biopsy punch (1 mm to 3 mm diameter) at Tokyo Medical and Dental University. The tissues were quickly submerged into RNAlater solution (Invitrogen) and then homogenized in Sepasol-RNA I Super G solution (Nakarai Tesque). Total RNA was isolated from tissue samples using the RNeasy Micro Kit (Qiagen, Germany) following the manufacturer’s instructions. Sequencing was performed on the NovaSeq6000 (Illumina) platform (150 bp paired end, 3.9 Gbp throughput on average).

### Quantification of gene expression

The raw reads were filtered by the following procedure: (i) removal of Illumina primers/adaptors/linker sequences; (ii) removal of terminal regions with continuous Phred-based quality<20 (48, 49). Reads that passed quality control were mapped to the mouse reference genome using STAR (version 2.7.9a) with the default parameter (50). After filtering ambiguous nucleotides and low-quality sequences, we quantified the expression of 21,826 coding genes in each sample. The unique read mapping rate against the reference mouse genome (GRCm39) ranged from 84.18 to 91.04% (Supplementary Table S1). Gene-level abundance estimation across samples was conducted using RSEM (51). The expected read counts from each sample were combined into a count matrix and were further normalized to transcripts per million (TPM) using in-house scripts. Both read count and TPM for each gene were used for downstream analyses.

### Identification of differentially expressed genes

The Pearson correlation matrix between sample-based gene expression profiles was calculated to deduce the similarity among biological replicates. The read count of each gene was normalized to counts per million (CPM). Genes with CPM < 2 were removed from downstream analysis. DESeq2 (52) was adopted to identify the differentially expressed genes (DEGs) between any pair stages. Genes with significant results (Benjamini–Hochberg adjusted p-value< 0.001; |log2 fold-change| > 1) were identified as DEGs in each comparison.

### Analysis of temporal expression patterns of DEGs

We used short time-series expression miner (STEM v1.3.12)(53) to investigate time-dependent expression profiles of DEGs with the following parameters: maximum unit change in model profiles between time points was 1; maximum output profile number was 20; the minimum ratio of the fold change of DEGs was no less than 2. A permutation-based test was used to determine the significance level of an identified transcriptome profile. Clusters with P < 0.05 were regarded as significantly enriched profiles.

### Establishment of expression profiles for auditory genes

To identify the expression profile of genes involved in hearing and inner ear development, we downloaded a list of auditory genes, which are known to be involved in deafness and/or the perception of sound in mice, from MGI (Mouse Genome Informatics). In addition, we included genes related to inner ear development and morphogenesis, including genes with GO annotation of “inner ear development,” “inner ear morphogenesis,” or “cochlea development.” We extracted DEGs according to this auditory gene lists for further analysis and visualization.

### Identification of temporal transcriptional regulatory networks

To further analyze the regulatory events among TFs and their targeted genes over time, we applied dynamic regulatory event miner (DREM2) (21) to reconstruct regulatory gene network during mouse inner ear development. We used mouse TFs and their targets provided by DREM2, including 336 TFs and 16,642 targeted genes. We filtered genes without prior TF interaction information and log-transformed the gene expression before the analysis, as recommended by the authors of DREM. We highlighted bifurcation events where the expression of a subset of genes diverges from the rest of the genes. Each bifurcation event is associated with a set of TFs that selectively regulate these events.

### Identification of the co-expression network

Weighted gene co-expression network analysis (WGCNA) using WGCNA 1.703 (54) was adapted to identify gene modules and gene co-expression networks associated with the inner ear developmental stages. Gene expression quantified by TPM was used as input. Pearson’s correlation matrix between all genes was calculated and further transformed into an adjacency matrix. With soft thresholding power by the “picksoftThreshold” function, we transformed the adjacency matrix into a topological overlap matrix (TOM) and identified modules using 1-TOM as the distance measure with a deepSplit value of 2. A minimum size of 30 for the resulting dendrogram and mergeCutHeight value of 0.25 were set to identify modules. A total of 143 genes that were unable to be assigned to any other module, were grouped into the module Grey. Pearson’s correlations between stage and module were calculated once the modules were identified. Finally, the co-expression networks were visualized using Cytoscape 3 (55).

### Enrichment analysis of Gene Ontology and KEGG

To understand the biological function/process of a set of genes, we performed the enrichment analysis of Gene Ontology (GO) (56) and KEGG pathways (57) using the R package *clusterProfiler* (58). P values were computed using Fisher’s exact test, and multiple test correction was performed using the Benjamini–Hochberg method (59) based on an FDR (false discovery rate) threshold of 0.05.

## Author contributions

J.L and D.K conceived, designed, and supervised the study. D.K performed the sampling. R.C analyzed and interpreted the data and drafted the manuscript. M.T, X.W, T.F, and T.N helped to interpret the data. All authors read and approved the final manuscript.

## Data accessibility

All raw data were stored in NCBI (BioProject: PRJNA823497). For further enquiry, please contact email

## Acknowledgement

We thank Ayumi Shimanuki for technically assisting the experiments.

