## Supplementary material for "Temporal and regulatory dynamics of the inner ear transcriptome during development in mice": Fig S1

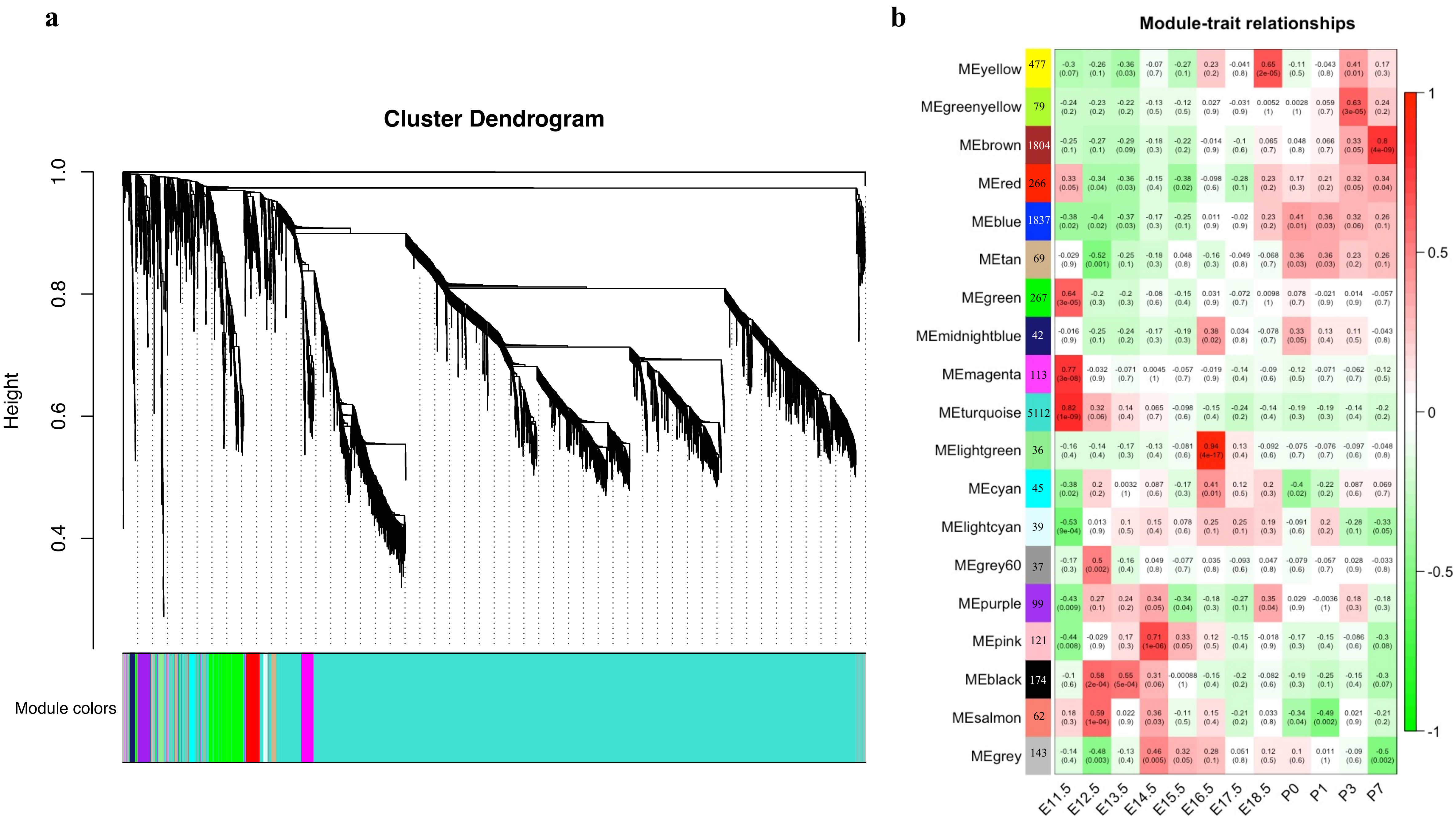


Figure S1. Co-expression network based on 10822 differentially expressed genes. (A) Hierarchical clustering of co-expression data. (B) Table of module–trait relationships. The value at the top of each square represents the correlation coefficient between the module eigengene and the trait with the correlation P-value in parentheses. The right panel is a color scale for module trait correlation from − 1 to 1.
